# Revealing Long-Term Multi-Factor Climate Impacts on Antarctic Phytoplankton: A Trend-Based Approach Using STL and Piecewise SEM

**DOI:** 10.1101/2025.06.03.657605

**Authors:** Hitomi Tanaka, Hideyuki Doi, Ryosuke Iritani

## Abstract

Climate change imposes multiple interacting stressors on ocean ecosystems. The Southern Ocean is among the regions where these impacts are particularly pronounced. Phytoplankton, which perform the bulk of primary production, both maintain the Antarctic marine food web and contribute to the role of the Southern Ocean as the largest oceanic sink for atmospheric CO_2_. Previous observational and modeling studies have revealed certain drivers–phytoplankton relationships and highlighted the important confounding factors. In this study, we quantified how climate and environmental drivers interact and influence phytoplankton biomass over a 20-year period, across the entire Southern Ocean, within a multi-factor framework. Specifically, we examined the relationships among atmospheric CO_2_, surface water pH, sea ice concentration, sea surface temperature, mixed layer depth, dissolved iron concentration, and chlorophyll *a* concentration. To perform this analysis, we introduced and applied two novel methodological approaches. Firstly, to address seasonal autocorrelation, we combined seasonal trend decomposition using LOESS (STL) with piecewise structural equation modeling (SEM). This approach allowed for robust estimation of long-term driver magnitudes and causal pathways, even in the presence of limited sample sizes and complex data structures. Secondly, given the highly interconnected nature of environmental drivers, we employed a data-driven SEM approach to objectively identify causal structures and avoid biases associated with a priori pathway selection. Our results show a clear signal of ocean acidification driven by increasing CO_2_, iron limitation of phytoplankton biomass during spring, and seasonal differences in both the magnitude and structure of the causal pathways, demonstrating the dynamic nature of seasonal processes in the Southern Ocean. This study presents the first Southern Ocean-wide, multi-factor assessment of climate and environmental impacts on phytoplankton biomass over a two-decade period. It introduces a methodological framework mitigates seasonal autocorrelation to accurately capture long-term seasonal trends, and demonstrates the utility of data-driven causal discovery in resolving complex ecological interactions.

## 1 Introduction

Climate change imposes multiple interacting stressors on ocean ecosystems [1], including warming [2], freshening [3][4], ocean acidification [5], sea-ice loss [6][7], increased water stratification [8], and altered nutrient regimes [9][10], among others. It is necessary to investigate the cumulative impacts of various concurrent stressors on marine systems[11]. The Southern Ocean is the largest oceanic sink for atmospheric CO_2_ [12] and is particularly sensitive to the impacts of climate change, including ocean acidification[13] and the melting of ice shelves and sea ice[14]. Given its major role in global carbon cycling and climate regulation, understanding the Southern Ocean’s response to these changes is essential for understanding the impact of climate change. Furthermore, the global ocean acts as a major carbon sink, having absorbed approximately 25–30% of anthropogenic CO_2_ emissions. Notably, about 40% of this uptake has occurred in the Southern Ocean [15]. Without this oceanic absorption, atmospheric CO_2_ concentrations would be roughly 50% higher than current levels [15]. This uptake of CO_2_ is partly driven by phytoplankton, which absorb CO_2_ through photosynthesis and transport it to the deep ocean via vertical fluxes [16]. Primary production by Southern Ocean phytoplankton also plays a key role in sustaining biodiversity and fueling the marine food web [17]. Therefore, studying phytoplankton dynamics in the Southern Ocean is essential for understanding the major impacts of climate change.

Recent modeling studies have advanced our understanding of the complex interactions among multiple environmental drivers and their influences on phytoplankton. Kwiatkowski et al.[18] projected simultaneous 21st-century ocean warming, acidification, deoxygenation, and reduced net primary production. Schmittner et al. [19] demonstrated that global nutrient and productivity patterns are highly sensitive to vertical mixing and organic-matter cycling. Fu et al. [20] showed that intensified stratification drives widespread declines in net primary and export production. Nevertheless, most of these studies still investigate some set of drivers, and multivariate assessments that compare the relative importance of several stressors remain scarce (e.g. [21]). Consequently, the approaches capable of identifying dominant factors and emergent interactions across a broader parameter space are still needed.

Here, we leverage two recent advances that make it feasible to assess the long-term, large-spatial scale, and multivariate interactions between climate forcing and ocean environmental variables. First, two decades of reanalysis and observation products (MERRA NOBM, COBE-SST, NSIDC, and others) provide monthly data on atmospheric CO_2_, surface water pH, sea ice concentration, sea surface temperature, mixed layer depth of the ocean, dissolved iron concentration and chlorophyll *a* concentration. Second, piecewise structural equation modeling (piecewise SEM) provides a robust framework to assess the relative importance of interacting environmental drivers and identify key causal factors[22][23]. It offers flexibility to accommodate non-normal distributions, temporal autocorrelation, and hierarchical structures commonly present in ecological data, while retaining the capacity to test causal hypotheses[24][23].

Using 240 monthly composites from 1998 to 2017, we ask:

1. Which environmental drivers exert the strongest direct and indirect controls on biomass of phytoplankton in the southern ocean?
2. Do the dominant pathways of the causal relationships among the variables vary seasonally?

By delivering the first basin-wide multivariate path analysis of the Southern Ocean, our study clarifies the long-term, large-spatial scale, and multivariate interactions between climate forcing and ocean environmental variables using the datasets.

However, conducting this analysis required addressing two remaining challenges: removing strong seasonal autocorrelation of the variables and developing a method to determine the underlying causal relationships.

The first challenge, seasonal autocorrelation, arises from the clear 12-month cycle present in the monthly time series (Fig. 1). If not properly accounted for, this seasonal pattern can distort the path coefficients and result in inaccurate conclusions when estimating long-term trends.

**Figure 1:**
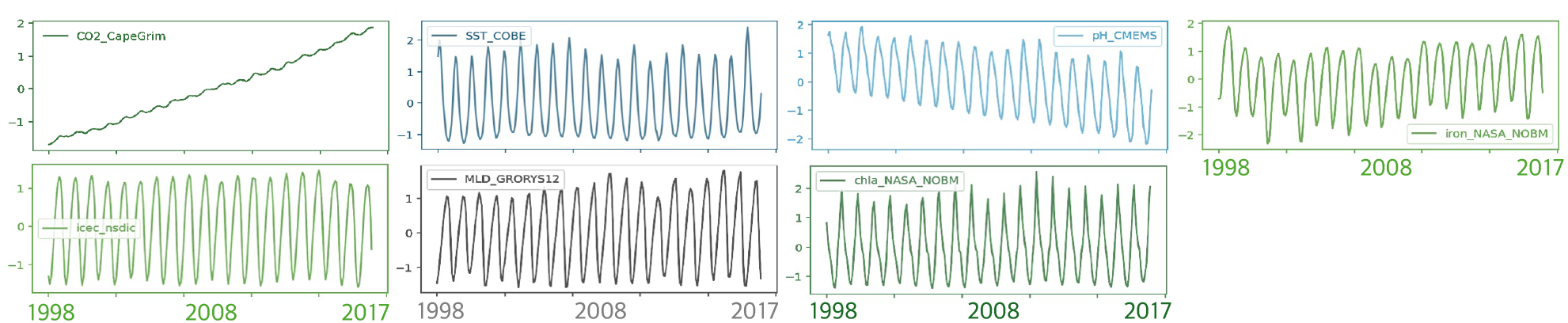
Original Dataset Exhibiting Strong Seasonal Autocorrelation Accurate estimation of long-term effects requires appropriately handling strong seasonal autocorrelation. If not addressed, this seasonality may affect the stability of path coefficients and lead to misleading conclusions about long-term trends.

Secondary, we face a key question: which path structure most accurately represents the system we aim to understand? SEM is attractive because it tests a hypothesized the network with global fit indices and provides the relative weight of every link [25]. To formulate a hypothesis about the path structure, previous studies have identified many pairwise relationships. For example, CO_2_ driven acidification [5][26], the impact of stratification on phytoplankton [9], and the ice albedo feedback that amplifies the warming of the sea surface [27]. Yet these effects form a densely interconnected web: rising CO_2_ warms the surface ocean, accelerates ice melt, and further amplifies surface temperatures through feedback processes. These changes influence phytoplankton, which in turn absorb CO_2_ through photosynthesis, triggering further interactions within the system.

When the network is this complex, deciding a priori which paths to include can bias the results, and the usual practice of adding or deleting links until the model fits may become arbitrary. Thus, although SEM remains a valid framework, a strictly hypothesis-driven version is impractical here.

### Addressing the Two Key Challenges

To overcome the two major challenges, namely seasonal autocorrelation and structural bias, we developed the following approach.

#### 1. Removal of Seasonal Autocorrelation

Accounting for or removing seasonal variation is occasionally done in ecological research when analyzing causal relationships [28][29]. As a method to take seasonality into account, classical seasonal autoregressive moving average models (SARMA) are well suited to capture seasonal dependence [30], but require relatively long records and cannot be embedded within the current implementation of piecewise SEM without substantial modification. Seasonal dummy variables offer a data efficient alternative [31], yet they average conditions within each season and therefore mask gradual, long-term change. Cubic spline or other flexible trend smoothers could, in principle, isolate long-term signals [31], but the resulting mixture of linear and higher-order polynomial terms is not currently compatible with piecewise SEM’s estimation engine.

We therefore implement Seasonal and Trend decomposition using Loess (STL) [32] as a preprocessing step. Removing the seasonal component suppresses the strong intra-annual autocorrelation while preserving the deseasonalized trend and anomaly series (Fig. 3). A piecewise SEM can then be fitted to the resulting data, with an AR(1) error structure added to the residuals [31]; this combination enables robust estimation of long-term driver magnitudes and pathways despite limited sample sizes and complex structure of data.

#### 2. Method to Determine the Underlying Structural Relationships without Prior Bias

To limit subjectivity, we conducted a data-driven SEM workflow. Data-driven causal discovery is an emerging tool for structuring the SEM models. Xu et al. [33] proposed a hybrid approach combining Bayesian networks and SEM, where a data-driven causal structure is first learned using a Bayesian network and then refined and validated with SEM, reducing reliance on first assumptions and improving model accuracy and reliability. The Bayesian networks also recover directed acyclic graphs like piecewise SEM[34], but their nonlinear probabilistic links are sometimes not compatible with the linear structure of SEM or piecewise SEM, and also it is difficult to take advantage of piecewise SEM’s strengths in handling non-normal data and hierarchical structures.

To address this, we implemented a beam search algorithm to explore candidate models that meet the goodness of fit index of piecewise SEM [24], and then selected the most appropriate model using a combination of information criteria[35][36] and prior ecological knowledge. This approach takes full advantage of the data structures that piecewise SEM can accommodate, while initially minimizing analyst bias by systematically exploring model space using goodness of fit indices and information criteria. At the same time, it allows for the selective incorporation of prior knowledge, achieving a balanced integration of data-driven discovery and expert-informed reasoning.

To clarify the multifaceted influence of increasing atmospheric CO_2_ on Antarctic phytoplankton, we first eliminate the pronounced seasonal cycle with STL [32]. We then analyzed the deseasonalized data using piecewise SEM with an AR(1) error term, a setup capable of handling limited sample sizes while accounting for residual autocorrelation [31]. In addition, we conduct a data-driven SEM framework which eliminate analyst bias. Our analyses provide a robust view of the long-term, basin-scale multivariate links among CO_2_ forcing, ocean environmental conditions, and phytoplankton dynamics.

## 2 Method

### Data

We used the publicly available datasets as follows and listed in Table 1.

- **Atmospheric CO**_2_**: CSIRO Cape Grim (time: 1976 - present**) [37] - Continuous Southern-Hemisphere background record with minimal urban or volcanic influence.
- **Sea surface temperature: COBE-SST v2 (time: 1850 - present, resolution: 1°)** [38] [39] - The longest 1° observational reconstruction, using an EOF-based statistical reconstruction with updated bucket corrections.
- **Sea ice concentration: NOAA/NSIDC Sea-Ice CDR v5 (1978 - present, 12.5 km)** [40] - Daily and monthly passive-microwave retrieval with cross-sensor calibration.
- **Surface water pH: Copernicus Surface Ocean Carbon Fields (1985 - present, monthly, 0.25°)** [41] [42][43]. - A neural-network reconstruction based on SOCAT ship observations, delivering gap-free global coverage.
- **Mixed-layer depth: Copernicus GLORYS12V1 (1993 - present, 1/12°)** [44][45] - Eddy-resolving physical reanalysis that assimilates T/S profiles, altimeter SSH, SST, and sea-ice. It provides finer resolution than ECCO (0.5°) or GODAS (1°) and supplies the 1990s record absent from Argo-based products.

**Table 1:**
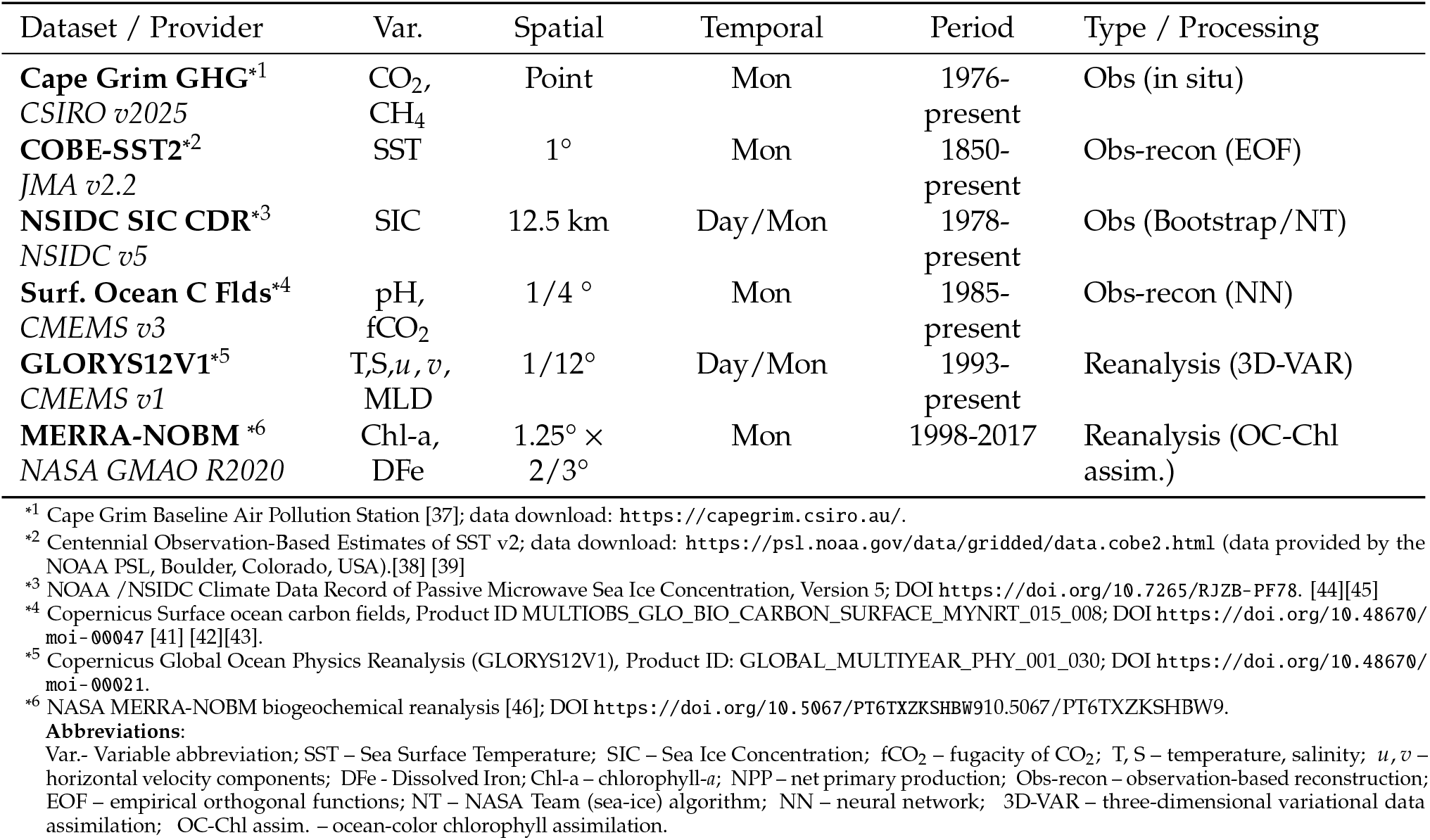
Summary of Datasets Used in This Study.

**Iron and chlorophyll *a*: NASA MERRA-NOBM (1998 - 2017, monthly, 1.25°** × **2/3°)** [46]-One of the few global biogeochemical reanalysis that simultaneously assimilates satellite chlorophyll and outputs iron together with chlorophyll in a fully consistent physical environment. It is the sole continuous, gap-free source for both variables.

These datasets were selected because they (i) provide at least 20 years of continuous data, (ii) minimize global spatial gaps, (iii) maintain internal physical-biogeochemical consistency, and (iv) are distributed through officially maintained portals with ongoing updates.

### Analytical workflow

The analytical workflow is shown in Fig 2, and the analysis was conducted through the following steps.

**Figure 2:**
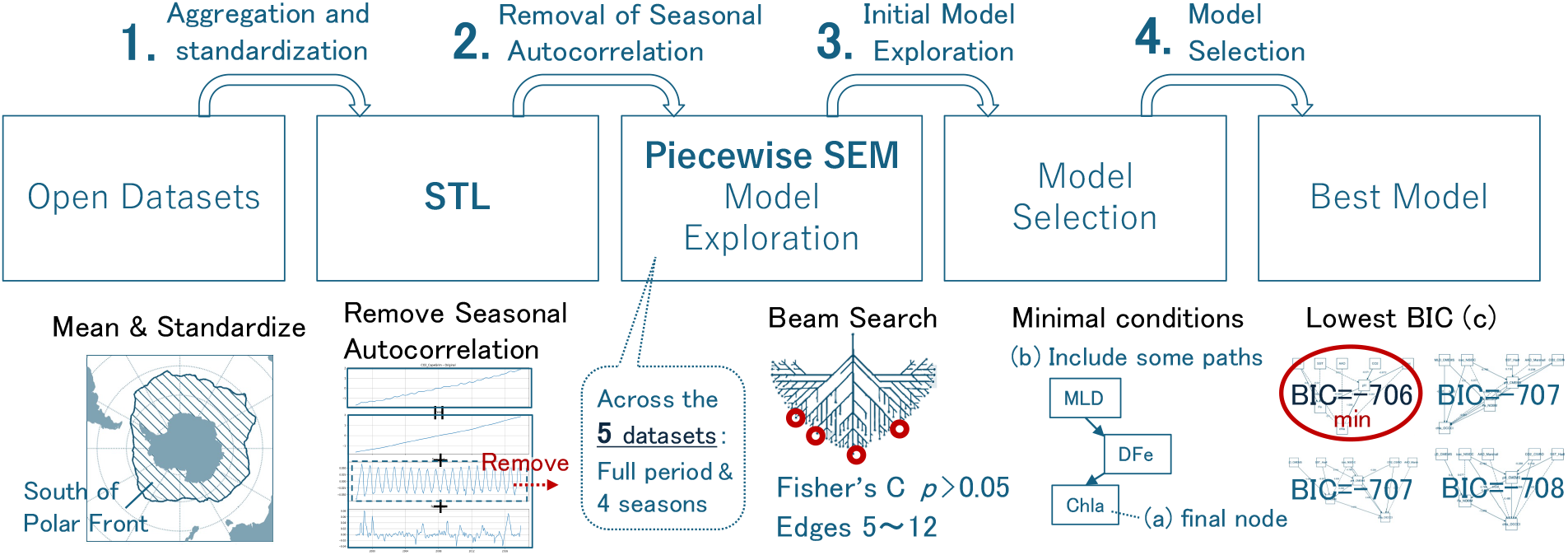
Analytical Workflow We applied latitude-weighted averaging and standardization to the datasets (1), removed seasonal autocorrelation of the variables using STL decomposition (2), used the beam search to explore the models satisfied the goodness-of-fit indices of piecewise SEM (3), and selected the best model with the lowest BIC after applying minimal conditions (4).

**Figure 3:**
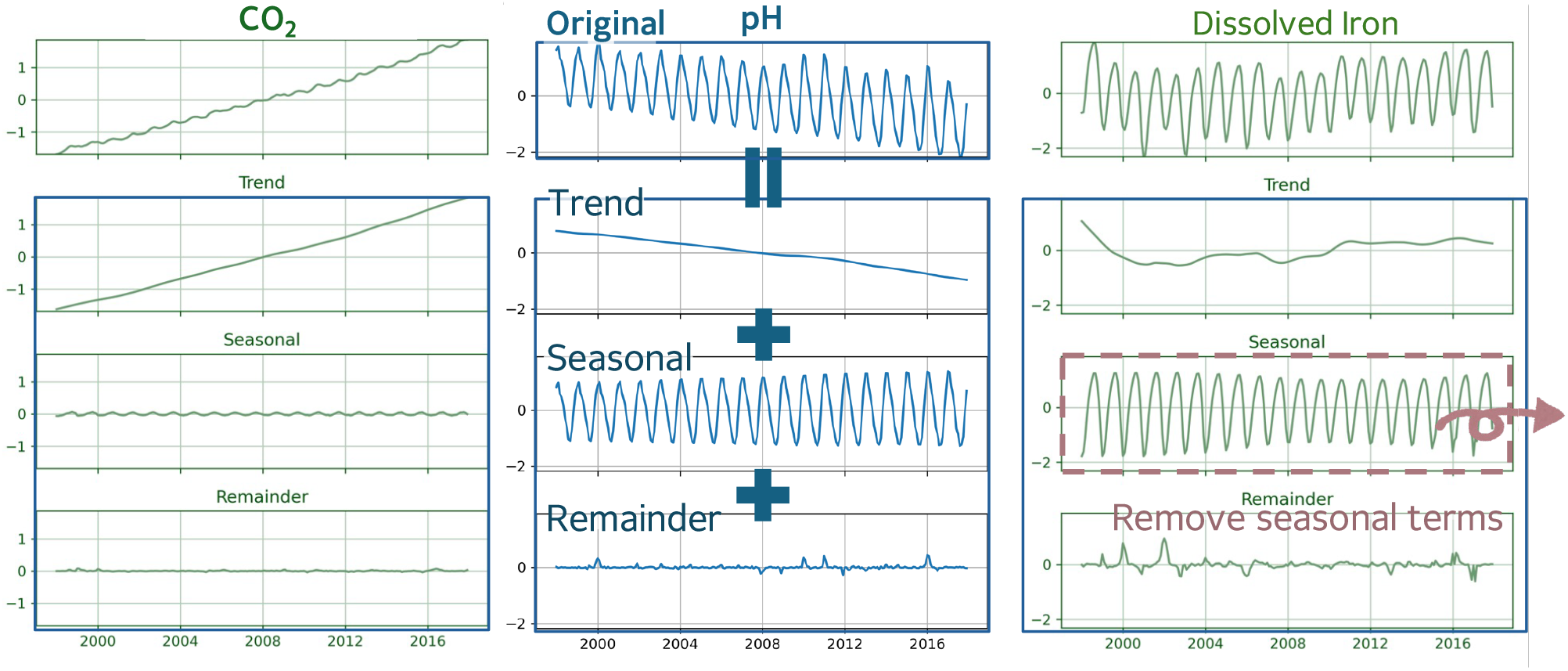
Examples of STL Applied Data STL is a method separating a time series into seasonal, trend, and remainder components using locally weighted regression. We applied STL to all variables and excluded the seasonal component, thereby facilitating more robust modeling and inference focused on underlying trends and residual variability.

#### 1. Aggregation and standardization

For every environmental parameter, the monthly means were calculated for all grid cells located south of the Polar Front (PF), the oceanic front that separates subantarctic water from Antarctic surface water. The south region of PF is known as a High Nutrient, Low Chlorophyll (HNLC) area where primary production is highly sensitive to climate change due to dissolved iron limitation and glacial melting. Since the PF is influenced by ocean currents and varies longitudinally, its position was determined by averaging data[47] from 2002 to 2014 at each longitude. Because a simple arithmetic mean of latitude-longitude gridded data overrepresents the higher latitudes, we first applied latitude-dependent area weighting and then computed the regional average. Each resulting time series was then standardized with the z-score.

#### 2. Removal of Seasonal Autocorrelation (Fig. 3, Details are provided in the following section.)

The standardized series were decomposed with Seasonal and Trend decomposition using Loess (STL). We removed the series with the seasonal component. The trend + remainder components are retained for subsequent analyses. Furthermore, after applying STL, the data were divided into seasonal periods: spring (September–November), summer (December–February), autumn (March–May), and winter (June–August). We then conducted the following analysis for five subsets: the full period and each individual season.

#### 3. Initial Model Exploration

Using the trend + remainder components series, we explored candidate path models within piecewise SEM framework across the five subsets, including the full period and each four individual season. A beam search algorithm enumerated models which satisfies Fisher’s C test (p_C_ > 0.05). The search space was constrained to models containing between 5 and 12 directed edges to limit overfitting and to avoid underfitting. Details on piecewise SEM are provided in the section after next.

#### 4. Model Selection

From the set of the models generated during the initial model explorationof piecewise SEM, we applied the minimal additional constraints based on the prior researches to refine the selection — (a) and (b) — and identified the best model accordingly (c). A summary is presented in Table 2.

**Table 2:** Applied Conditions and Number of Candidate Models for Each Season in 4. Model Selection.

| Season | Model counts from (a) | Applied conditions | Possible combinations | Model counts meeting applied conditions |
| --- | --- | --- | --- | --- |
| Spring | 25150 | (a) | $1.1 \times 10^{17}$ | 25150 |
| Summer | 968 | (a) & (b-2) | $2.7 \times 10^{14}$ | 28 |
| Autumn | 7385 | (a) & (b-1) | $1.4 \times 10^{13}$ | 443 |
| Winter | 3 | (a) | $1.1 \times 10^{17}$ | 3 |
| Full Period | 9347 | (a) & (b-1) | $1.4 \times 10^{13}$ | 126 |

a. We retained only those models in which Chl-*a* appeared exclusively as the terminal node of the partial order. The number of possible path combinations is theoretically estimated at 1.1 × 10^18^ (for details on the combinatorial calculation, see Fig. S1). For spring and winter, step (b) was skipped, and the analysis proceeded directly to step (c) after applying this condition.
b. If the number of candidate models was still too large to manage in (a), each model was required to include three literature supported “white list” pathways.
  (b-1) We included pathways identified in previous studies, such as the effect of increased CO_2_ on sea surface temperature (CO_2_ → SST) and the influence of mixed layer depth on dissolved iron and subsequently on chlorophyll-a concentration (MLD → DF e → Chl-a). This was applied to the Full Period and autumn analyses. The number of possible path combinations is 1.4 × 10^13^ (Fig. S1).
  (b-2) For summer, none of the previously identified MLD → DFe pathways were detected. Therefore, we included only the CO_2_ → SST and DFe → Chl-a paths from step (b-1). The number of possible path combinations is 2.7 × 10^14^ (Fig. S1).
c. From the remaining set, the model with the lowest Baysian Information Criterion (BIC) was chosen as the final model. BIC is a model selection metric that balances model fit and complexity by penalizing the number of parameters, and is designed to consistently select the true model among candidates.

### STL and Autocorrelation Reduction

To address seasonal variation, we applied Seasonal and Trend decomposition using Loess (STL) [32]. STL decomposes a time series (*Y*_*t*_) into three additive components: trend (*T*_*t*_), seasonal (*S*_*t*_), and remainder (*R*_*t*_), as expressed by:

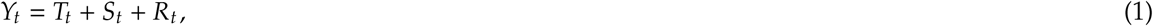

STL uses LOESS, a local regression technique, to iteratively estimate *T*_*t*_ and *S*_*t*_ in a flexible and robust manner.

After estimating and removing the seasonal component, we obtain a seasonally adjusted series:

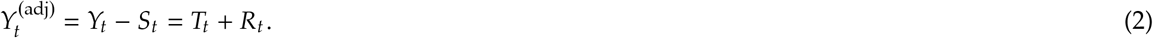

This adjustment is crucial because strong seasonal patterns often introduce significant autocorrelation in the data. By removing *S*_*t*_, the resulting series 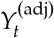 shows reduced autocorrelation, allowing for more effective modeling and inference based on residual dynamics and trend. Examples of data after STL decomposition are presented in Fig. 3.

We performed STL using the `statsmodels` package in Python. The Python version used was 3.12.8, and the `statsmodels` [48] version was 0.14.4. The code is shown below.

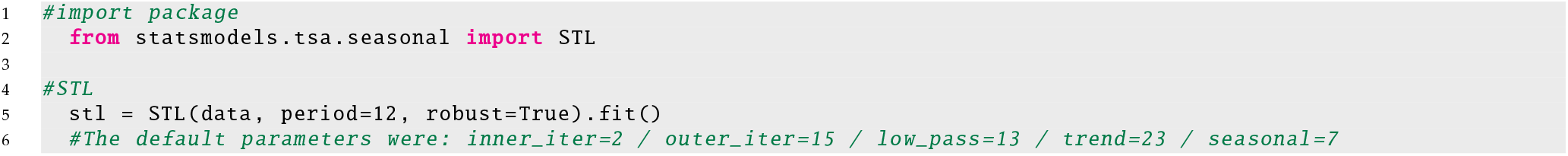

The variable `data` refers to the time series data for each climate variable. `period = 12` indicates the periodicity of the sequence, in this case corresponding to 12 months. Setting `robust = TRUE` enables the algorithm to account for outliers. Regarding the default parameters: `inner_iter = 2` ensures greater stability in the separation of seasonal and trend components, though theoretically one iteration is sufficient. `outer_iter = 15` performs 15 rounds of robustness iterations to down-weight outliers; this can be reduced to 10 if outliers are few, to save computation time. `low_pass = 13` is the smallest odd number greater than the period (12), effectively filtering out residual high-frequency noise. `The combination trend = 23` and `seasonal = 7` follows the recommendation of Cleveland [32], and produces a suitably smooth trend component for monthly data.

### Piecewise SEM

Piecewise Structural Equation Modeling (SEM) estimates each path as an independent generalized (mixed) model and links them through the rules of graphical d-separation [22]. The results of basis set of conditional independence claims provides a global goodness-of-fit test via Fisher’s C statistic [24].

Since parameters are obtained from separate regressions rather than a single covariance matrix, piecewise SEM needs fewer observations than conventional covariance based SEM while still delivering unbiased path coefficients and valid standard errors.

The framework is inherently flexible: any Generalized Linear Model (GLM)/ Linear Mixed Model (LMM)/ Generalized Linear Mixed Model (GLMM) can be inserted into a path, enabling “generalized multilevel path models” that mix link functions, random effects, and distribution families [49]. Temporal or spatial autocorrelation can be handled by specifying residual structures such as AR(1) within each component model.

Thus, piecewise SEM lets the researchers evaluate complex causal networks with modest data and realistic error terms, broadening the applicability of path analysis across disciplines.

We applied Generalized Least Squares (GLS) to the STL and autocorrelation adjusted data *Y*^(adj)^, incorporating an AR(1) structure in the residuals to account for temporal autocorrelation by modeling each month’s residual as dependent on the previous month’s, thereby capturing the persistence commonly observed in monthly time series data. The model was estimated in R using equations such as the following. The R version used was 4.4.3, with the `piecewiseSEM` package [49] version 2.3.0.1 and the `nlme` package [50][51] version 3.1-166.

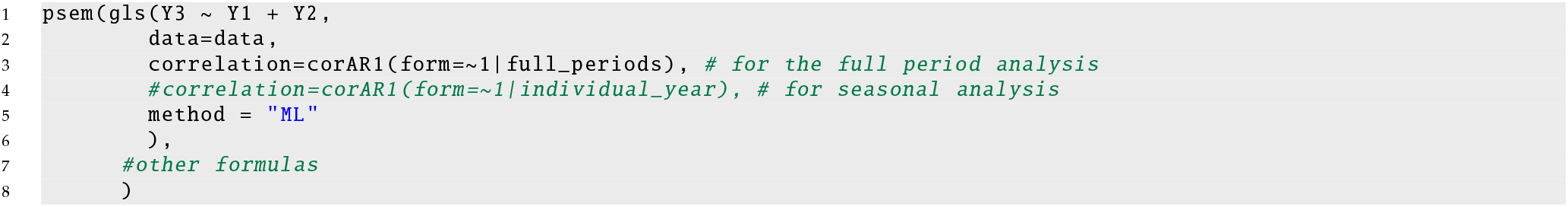

Here, Y1, Y2 and Y3 represent example variables, such as CO_2_ and Chl *a*, corresponding to the autocorrelation-adjusted time series 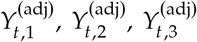 introduced in the previous section. The `corAR1` specifies an AR(1) structure in the residuals, applied to the full time series period referred to as the Full Period analysis. While in the seasonal analyses, `corAR1` was applied only within individual years, since, for example, late spring in one year is not temporally correlated with early spring in the following year. The argument `method = “ML”` specifies that model estimation was performed using maximum likelihood (ML), which is required when comparing models using information criteria such as BIC [31].

## 3 Results

The piecewise-SEM results for the full period and each season are presented in Fig. 4, with global-goodness of fit indices and information criteria summarized in Table 3. The following subsections provide a detailed description of the results for each season and the full-period analysis.

**Table 3:** Values of Global Goodness of Fit Indices and Information Criteria.

| Season | Global-goodness of fit indices |  | Information criteria |  |
| --- | --- | --- | --- | --- |
|  | Ficher's C | Ficher's C p-value | BIC | AIC |
| Full Period | 30.921 | 0.056 | -2708.733 | -2809.672 |
| Spring | 23.294 | 0.385 | -706.357 | -764.998 |
| Summer | 24.498 | 0.139 | -229.302 | -347.923 |
| Autumn | 19.198 | 0.509 | -805.553 | -866.289 |
| Winter | 25.636 | 0.178 | -896.299 | -957.035 |

**Figure 4:**
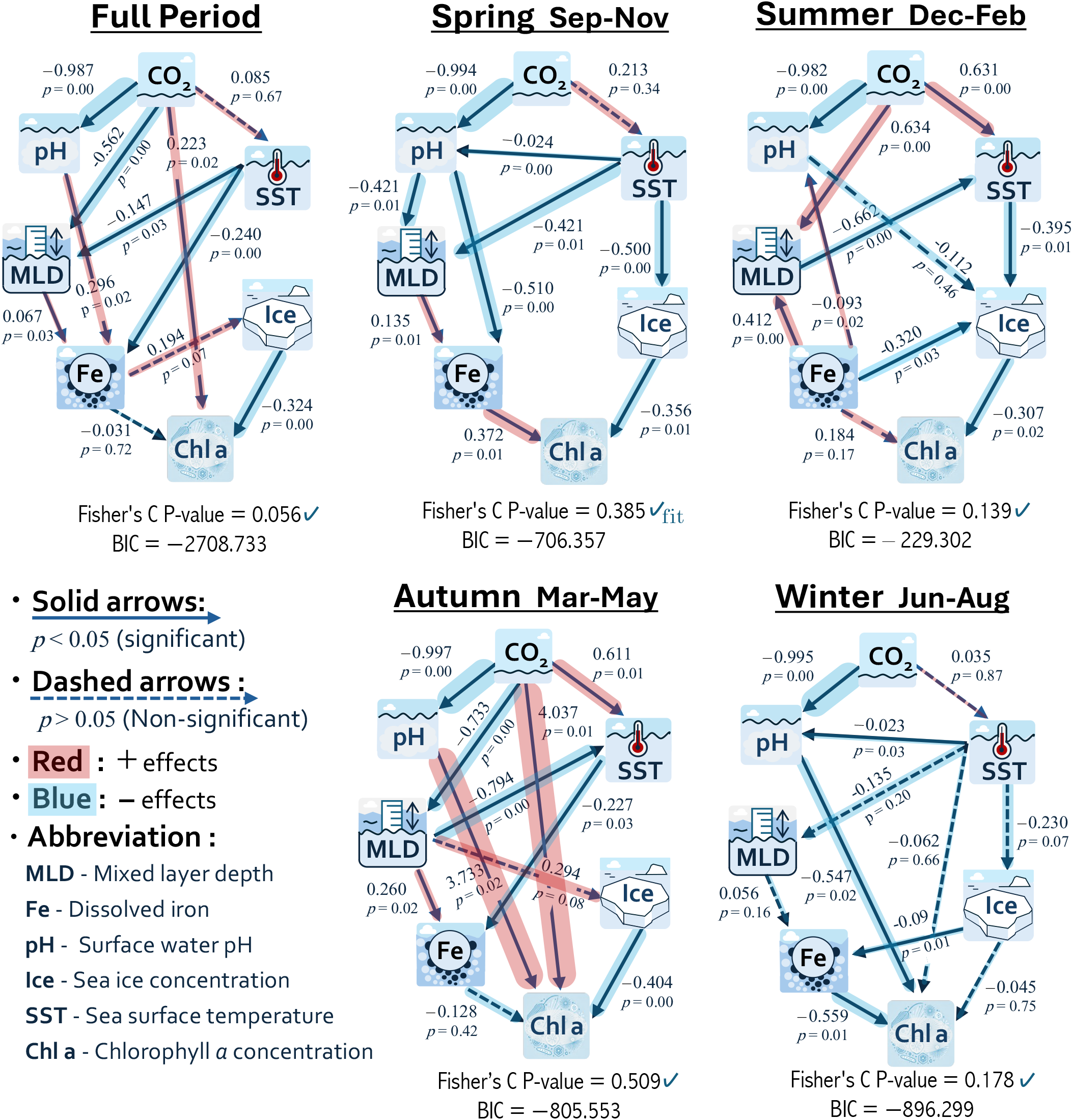
Piecewise SEM Results (*R*^2^) of the Best Models for the Full Period and Individual Seasons The figures are the results of the best model based on the lowest BIC and validated by Fisher’s C test. Solid arrows indicate statistically significant paths (*p <* 0.05) and are the primary focus of interpretation. Red highlights represent the positive effects, blue highlights indicate negative effects, and thicker highlights correspond to larger path coefficients. The figures show both direct and indirect causal relationships among variables; the magnitude of each path coefficient reflects the strength of the relationship.

### Spring

An increase in CO_2_ leads to a decrease in surface water pH, indicating that ocean acidification appears as the most influential pathway. Only in spring, an increase of sea surface temperature results in a decrease in sea ice concentration, clearly reflecting the reduction of sea ice due to warming. Furthermore, in spring, higher iron concentrations are associated with increased chlorophyll *a*, suggesting that iron limitation, a characteristic of the Southern Ocean as a high nutrient low chlorophyll (HNLC) region, becomes particularly prominent during this season. There is also a negative relationship between sea ice concentration and chlorophyll *a*, likely because phytoplankton growth is restricted to limited the open water areas during periods of extensive sea ice coverage.

### Summer

An increase in CO_2_ consistently lowers pH, underscoring ocean acidification as the dominant pathway. The results also show a clear signal of ocean acidification driven by the increase in CO_2_ contributes to higher sea surface temperature, and that an increase in SST is associated with a reduction in sea ice extent.

### Autumn

In autumn as well, an increase in CO_2_ leads to a decrease in pH, indicating that ocean acidification remains the influential pathway. CO_2_ is also found to have a direct effect on increasing sea surface temperature.During this season, all parameters show relatively strong effects, with particularly strong relationships observed between CO_2_, pH, and chlorophyll *a*.

### Winter

In winter as well, an increase in CO_2_ leads to a decrease in pH, indicating that ocean acidification remains the most dominant pathway. Compared to the other seasons, the influence of almost all other pathways is relatively weak. This may be attributed to the reduced ocean dynamics during the winter, which limits the expression of the interactions among the variables as distinct pathways.

### Full Period and Seasonal Comparisons

Across all seasons, an increasing of CO_2_ consistently leads to a decrease in pH, indicating that ocean acidification is a dominant pathway, with path coefficients close to 1. The relationship between deeper mixed layer depth and higher iron concentrations, suggesting the greater nutrient availability, appears clearly in spring, and to a lesser extent when the full period are considered together. Although increasing CO_2_ is generally associated with global surface warming through the greenhouse effect, a clear linear relationship between CO_2_ and sea surface temperature was observed only in summer and autumn in the Southern Ocean. An increasing of sea surface temperature was associated with a shallower mixed layer depth in both spring and the full period analysis, suggesting the increasing of surface temperature increases stratification. In autumn, the relationship appeared reversed, but overall, a negative correlation between sea surface temperature and mixed layer depth was evident. Furthermore, the path exploration conducted separately for the full period and individual seasons revealed notable differences not only in the magnitude of pathways but also in their structural relationships. These findings highlight that the Southern Ocean exhibits distinct seasonal dynamics.

## 4 Discussion

In this study, we analyzed the impact of rising atmospheric CO_2_ and multiple oceanic drivers on phytoplankton in the Southern Ocean south of the Polar Front, over a 20-year period from 1998 to 2017. With removing seasonal variation of the variables, we firstly applied Seasonal and Trend decomposition using Loess (STL) for piecewise SEM to capture complex relationships among key ecological variables. Using a data-driven approach, we derived a wide range of insights that extend beyond the explanatory scope of traditional hypothesis-driven SEM.This discussion is structured as follows: we first highlight the key findings, then provide considerations on the methodological approaches employed, and finally present the study’s conclusions.

### Key Findings

We revealed a range of important findings regarding the biogeochemical and ecological responses of the Southern Ocean to phytoplankton chlorophyll *a*. In the following, we summarize key insights related to ocean acidification, iron limitation, and seasonal differences in these processes.

1. The consistent reduction in pH with increasing CO_2_ across all seasons, identifying ocean acidification as the dominant pathway. Recent studies have highlighted ongoing ocean acidification in the Southern Ocean. Zemskova et al.[52] showed that dissolved inorganic carbon decreased in the 1990s and 2000s but has increased in surface waters since the 2010s south of 60 Metzl et al. [53] reported a long-term decline in annual mean surface pH from 1985 to 2020 south of the Antarctic Convergence, confirming sustained acidification. This study demonstrates that ocean acidification is not only ongoing but pronounced across the entire Southern Ocean south of the Polar Front over multi-decadal timescales.
2. In spring, elevated iron concentrations correlate with higher chlorophyll *a* levels, indicating that iron limitation in the Southern Ocean is particularly significant during this season. Observational studies across the Southern Ocean have consistently shown that iron is a key limiting factor for phytoplankton growth[54] [55] [56]. Satellite data show large blooms near iron sources, and models reveal that iron supply influences plankton composition and export flux—together supporting Martin’s iron hypothesis and decades of Southern Ocean research[57]. Our result provides the first large-spactial scale, long-term causal analysis confirming dissolved iron limitation for phytoplankton production across the Southern Ocean, with a novel focus on springtime dynamics using observation, reanalysis data, and path modeling.
3. Distinct seasonal differences were observed in both the magnitude and structure of the causal pathways, demonstrating the dynamic nature of seasonal processes in the Southern Ocean. The previous studies have also highlighted various seasonal variations in the Southern Ocean. For example, Tagliabue et al. tagliabue2014surface show that deep water mixing during winter replenishes the mixed layer with iron from subsurface reservoirs. In spring, this iron is rapidly depleted, and summer, vertical iron supply becomes minimal. During this stratified period, phytoplankton increasingly rely on recycled iron, highlighting the importance of internal cycling processes until deep mixing resumes in autumn. Our results also reveal strong seasonal variation in the relationships among mixed layer depth, dissolved iron, and Chl a. However, when viewed over the Full Period, these relationships appear to have a relatively minor overall influence.

While these findings offer valuable insights into oceanic influences on phytoplankton, a more comprehensive understanding would require the inclusion of atmospheric climate variables as well. Our analysis targeted the ocean’s response to rising atmospheric CO_2_, but other atmospheric forcings still warrant explicit treatment. Variations in stratospheric ozone, aerosol loading, photosynthetically-active radiation (PAR), and the Antarctic Oscillation jointly reshape UV exposure [58], winds, and nutrient supply [59], and could therefore alter the pathways we resolved. Integrating these drivers into future structural models will be essential for a complete causal picture. Also, our analysis integrated datasets from multiple sources, each with its own uncertainty characteristics. While we selected variables known to perform reliably in polar regions, uncertainties remain. Continued data collection and reanalysis of biogeochemical fields will help reduce these uncertainties and are expected to further advance understanding of the Southern Ocean system.

### Considerations on the Methodological Approach Adopted in This Study

To remove seasonal autocorrelation, we applied STL [32] to remove seasonal effects, enabling piecewise SEM analysis with an AR(1) residual structure [31]. This approach allows us the reliable estimation of long-term driver effects and causal pathways, with limited sample sizes and complex data structures. Although the previous ecological studies have used deseasonalized residuals or anomalies for statistical analyses (e.g., [28] [29]), and the others have incorporated seasonal indices directly into SEMs (e.g. [60]), To our knowledge, there are no evidence of STL-filtered data being directly used in SEM or piecewise SEM. Our approach with removing seasonal components via STL and analyzing the resulting trend and anomalies with piecewise SEM, offers anovel contribution, particularly in long-term dataset with seasonal dynamics,where such applications remain largely unexplored.

As for data-driven SEM, we applied a beam search strategy to identify the candidate models that satisfy the goodness-of-fit requirements of piecewise SEM, and subsequently refined model selection by integrating information theoretic metrics with relevant ecological insights. Data-driven causal discovery is emerging as a promising alternative to traditional correlation analysis [33][61], allowing the data itself, rather than relying on prior assumptions, to identify the most plausible pathways. However, its application to Southern Ocean phytoplankton remains unexplored, making this a novel approach in the context of this study. Our data-driven method leverages the flexibility of piecewise SEM to handle complex data structures, while reducing analyst bias in the early stages by using a systematic search based on fit statistics and information-theoretic criteria. It also enables the targeted use of prior ecological knowledge, offering a balanced framework that combines data-driven exploration with theoretical grounding. While traditional research typically tests predefined hypotheses, in this study, we take a data-driven approach that integrates empirical patterns with theoretical insights to uncover underlying causal structures. The data-driven SEM method is powerful in extracting information beyond what hypothesis-driven models can offer, but its flexibility also requires careful handling to avoid subjective bias. The model space under our constraints included at least 10^13^ possible structures(see S1 4.), indicating the data-driven foundation of the approach. Importantly, to generalize this method, future research must establish transparent reporting standards, including the size of the model space, the number of models evaluated, and the rationale for constraints, and promote reproducibility through open data and clear workflows.

## Conclusion

This study conducted a comprehensive, multi-factor structural analysis of phytoplankton biomass in the Southern Ocean and key ocean environmental variables using an unprecedented 20-year record (1998–2017) spanning the entire Southern Ocean south of the Polar Front. We identified a clear signal of ocean acidification driven by the increasing of CO_2_, persistent iron limitation of phytoplankton, and a pronounced intensification of this iron stress during spring. We also found that the strength and even the direction of many relationships varies markedly with season. To analyze the long-term impacts of complex drivers with strong seasonal autocorrelation, we developed a novel framework combining STL with piecewise SEM. By applying SEM in a data-driven way and interpreting the results alongside existing knowledge, we were able to uncover new insights into the impact of increasing CO_2_ and multiple oceanic drivers on phytoplankton in the Southern Ocean. While this study focused on atmospheric CO_2_ and oceanic variables to phytoplankton biomass, future work can incorporate broader climate drivers, such as stratospheric ozone, the Antarctic Oscillation, aerosols, and photosynthetically active radiation. Including these factors will help build a more integrated causal picture of the Southern Ocean system.

## Acknowledgements

This study was conducted using the datasets (i) Cape Grim Greenhouse Gas Monitoring Data v2025-02 (CSIRO Environment & Australian Bureau of Meteorology); (ii) Surface Ocean Carbon Fields (DOI `10.48670/moi-00047`) and (iii) Global Ocean Physics Reanalysis (DOI `10.48670/moi-00021`) as EU Copernicus Marine Service; (iv) NASA GMAO MERRA-NOBM monthly reanalysis R2020; (v) NOAA/NSIDC passive-microwave sea-ice concentration CDR v5 (DOI `10.7265/rjzb-pf78`); and (vi) COBE-SST2 SST & sea-ice v2 (data provided by the NOAA PSL, Boulder, Colorado, USA, from their website at (https://psl.noaa.gov). CSIRO and the Australian Bureau of Meteorology make no warranty, expressed or implied, regarding the Cape Grim data. The authors thank JSPS KAKENHI for funding (22K18429 and 22K12461 to HD; 24H01528 and 24H02291 to RI).

## Supplementary Material

**Figure S1:**
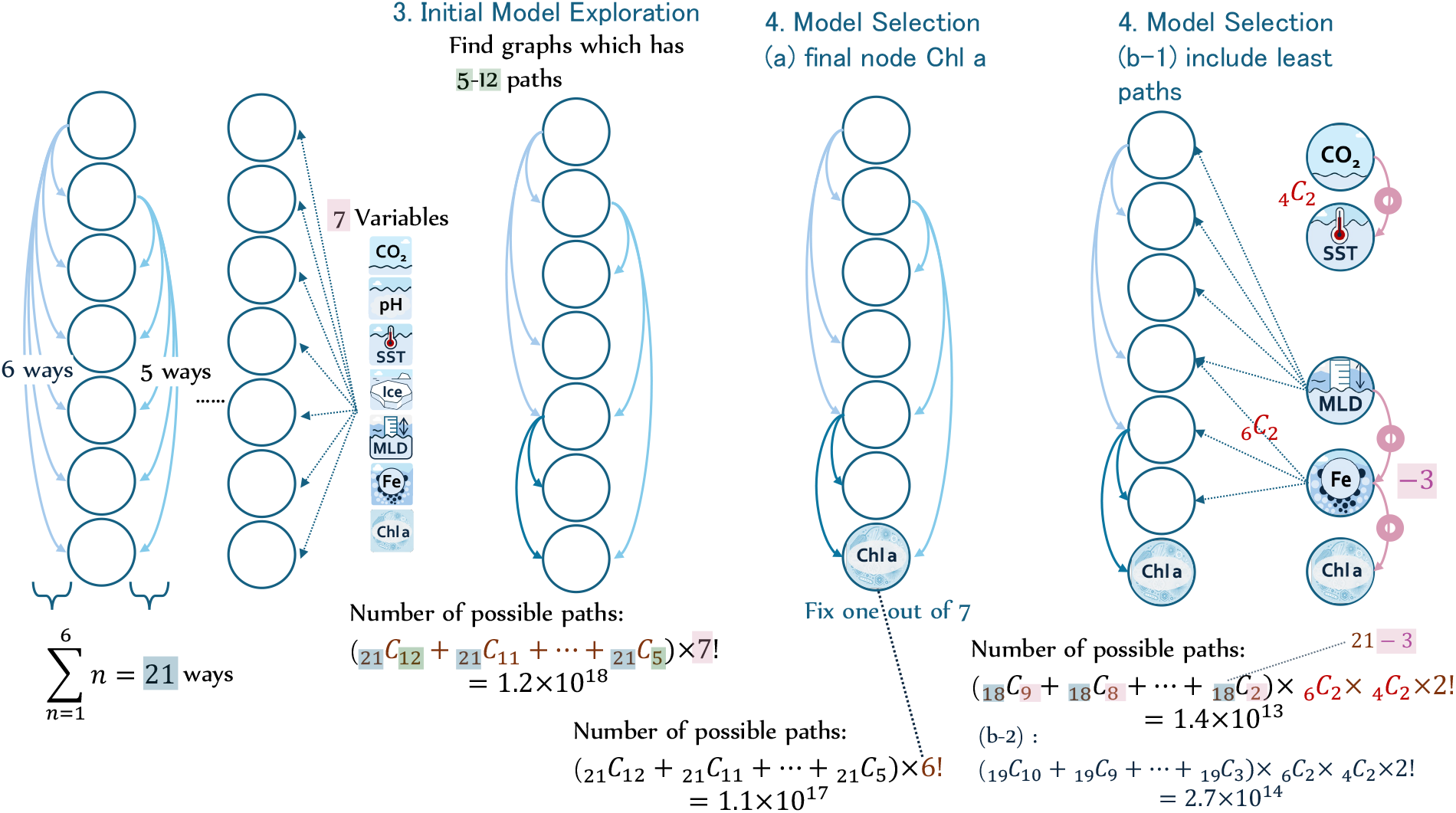
Number of possible paths

